# Metabolic labeling of RNAs uncovers hidden features and dynamics of the *Arabidopsis thaliana* transcriptome

**DOI:** 10.1101/588780

**Authors:** Emese Xochitl Szabo, Philipp Reichert, Marie-Kristin Lehniger, Marilena Ohmer, Marcella de Francisco Amorim, Udo Gowik, Christian Schmitz-Linneweber, Sascha Laubinger

**Author notes:** These authors contributed equally (Emese Xochitl Szabo, Philipp Reichert). Corresponding author: Sascha Laubinger.

## Abstract

Transcriptome analysis by RNA sequencing (RNA-seq) has become an indispensable core research tool in modern plant biology. Virtually all RNA-seq studies provide a snapshot of the steady-state transcriptome, which contains valuable information about RNA populations at a given time, but lacks information about the dynamics of RNA synthesis and degradation. Only a few specialized sequencing techniques, such as global run-on sequencing (GRO-seq), have been applied in plants and provide information about RNA synthesis rates. Here, we demonstrate that RNA labeling with a modified, non-toxic uridine analog, 5-ethynyl uridine (5-EU), in *Arabidopsis thaliana* seedlings provides insight into the dynamic nature of a plant transcriptome. Pulse-labeling with 5-EU allowed the detection and analysis of nascent and unstable RNAs, of RNA processing intermediates generated by splicing, and of chloroplast RNAs. We also conducted pulse-chase experiments with 5-EU, which allowed us to determine RNA stabilities without the need for chemical inhibition of transcription using compounds such as actinomycin and cordycepin. Genome-wide analysis of RNA stabilities by 5-EU pulse-chase experiments revealed that this inhibitor-free RNA stability measurement results in RNA half-lives much shorter than those reported after chemical inhibition of transcription. In summary, our results show that the Arabidopsis nascent transcriptome contains unstable RNAs and RNA processing intermediates, and suggest that half-lives of plant RNAs are largely overestimated. Our results lay the ground for an easy and affordable nascent transcriptome analysis and inhibitor-free analysis of RNA stabilities in plants.

## Introduction

Transcriptomes are highly dynamic and vary greatly among different cell types, environmental conditions and developmental stages. Plant transcriptome analysis by RNA sequencing (RNA-seq) has become an affordable key tool for understanding plant development and adaptive growth, for assessing the effects of specific mutations, and for genome annotation. Different types of RNA-seq technologies have not only been used for transcript abundance measurements in plants, but also for the detection and annotation of transcript sequence variation generated by alternative splicing, differential promoter usage or by the selection of different transcription termination and polyadenylation sites (Sherstnev et al., 2012; Morton et al., 2014; Filichkin et al., 2015; Kappel et al., 2015; Abdel-Ghany et al., 2016; Cartolano et al., 2016; Sun et al., 2017; Tokizawa et al., 2017; Ushijima et al., 2017; Vaneechoutte et al., 2017; Zhang et al., 2017). RNA-seq analysis of mutants impaired in RNA turnover or the enrichment of specific RNA fractions prior to library preparation also provided important information about the nature, the stability and the processing of various RNA molecules as well as their modifications (Drechsel et al., 2013; Schwarzl et al., 2015; Vandivier et al., 2015; Li et al., 2016; Shen et al., 2016; Cui et al., 2017; David et al., 2017; Sorenson et al., 2018).

Most transcriptomic studies in plants focus on the detection of steady-state transcriptomes that is defined by the rate of RNA synthesis and degradation. Although the steady-state transcriptome provides essential information about the RNA composition, information about RNA synthesis and degradation cannot be deduced from steady-state transcriptomes. Furthermore, the presence of long-lived and abundant RNAs in steady-state transcriptomes limits the detection of low abundant and unstable RNAs including RNA processing intermediates. Certain techniques take such challenges into account. Among them, global run-on sequencing (GRO-seq) has been applied in plants (Erhard et al., 2015; Hetzel et al., 2016; Liu et al., 2018). For GRO-seq analyses, labeled nucleotides such as bromouridine (BrU) are added to isolated nuclei. Transcriptionally engaged RNA polymerase II completes the transcription of its bound genes and incorporates BrU into the nascent RNAs (Lopes et al., 2017). BrU allows affinity purification of labeled RNAs with antibodies, followed by sequencing analysis. GRO-seq facilitates detection of transcriptionally active genes and allows to disentangle the relative contributions of RNA synthesis and degradation when compared to the steady-state transcriptome. GRO-seq is, however, an *in vitro* technique, which limits its applications including the expression analysis under different environmental conditions. Furthermore, RNA processing and degradation cannot be studied in standard GRO-seq experiments. Another technique that specifically enriches nascent RNA molecules is native elongating transcript sequencing (Net-seq) (Churchman and Weissman, 2011). For Net-seq analysis, engaged RNA polymerase complexes are immunopurified followed by isolation of nascent RNA molecules from the RNA-DNA-polymerase ternary complex. This method permits estimations of polymerase dynamics and RNA processing (Mayer et al., 2015; Nojima et al., 2015; Harlen et al., 2016; Mayer and Churchman, 2016; Zhu et al., 2018). Net-seq also allows the detection of unstable transcripts that escape detection using conventional RNA-seq (Marquardt et al., 2014; Wery et al., 2018).

Another set of techniques for studying transcriptome dynamics and homeostasis employs metabolic RNA labeling. This approach provides information about neo-synthesized RNAs during the labeling period including nascent RNAs, unstable RNA species and RNA processing intermediates (Rabani et al., 2011a). For metabolic labeling of RNAs, cells are incubated with nucleotide analogs such as BrU, 4-thiouridine (4-SU) or 5-ethynyl uridine (5-EU). These analogs are incorporated during transcription instead of unlabeled uridine. Labeled RNAs can be purified by specific antibodies or by adding a biotin tag to the RNA molecule by click-chemistry followed by streptavidin affinity purification (Jao and Salic, 2008; Duffy et al., 2015; Yildirim, 2015; Sawant et al., 2016; Herzog et al., 2017). Such pulse-labeling experiments using 4-SU combined with DNA array hybridization revealed RNA synthesis rates in Arabidopsis (Sidaway-Lee et al., 2014). Labeling of nucleolar RNAs with 5-EU has been applied to visualize plant nucleoli (Dvorackova and Fajkus, 2018). Although RNA labeling with base analogs occurs in living cells, its current application is mostly limited to cell cultures and, additionally, some base analogs exhibit toxic effects (Burger et al., 2013). DNA-containing organelles pose an additional problem for metabolic labeling of RNA given their additional membrane barriers. Still, using 4-SU or 5-EU, successful labeling of RNAs has been reported for mitochondria in human cell lines (Borowski and Szczesny, 2014; Nguyen et al., 2018). By contrast, RNA labeling using modified bases has not yet been reported for plant organelles.

Metabolic labeling of RNAs also allows determination of RNA stabilities. Traditionally, RNA half-lives are determined by following RNA degradation over time after blocking transcription by specific molecules. These experiments employ cordycepin, α-amanitin or actinomycin D to inhibit RNA synthesis. The disadvantages of these methods are that the specific chemicals are often toxic and that they do not exert their full inhibitory capacity immediately after the treatment, causing a lag phase before transcription is blocked efficiently (Wada and Becskei, 2017). Transcription and RNA degradation is also tightly linked (Wada and Becskei, 2017). Thus, blocking transcription in turn can positively affect RNA stability and biases RNA half-live calculations (Braun and Young, 2014). Inhibitor-free strategies for RNA stability measurements, such as 5’Bromouridine IP Chase (BRIC)-seq, start with pulse labeling using a nucleotide analog followed by chasing periods of various lengths using unlabeled nucleotides (Tani et al., 2012b; Imamachi et al., 2014). Transcriptome analysis after chasing periods of different length provides information about changes in RNA abundance over time. RNA half-lives determined by pulse-chase experiments often resulted in different stability measurements compared to inhibitor-based approaches (Rabani et al., 2011b; Tani et al., 2012b).

Here, we present and propose the application of 5-ethynyl uridine (5-EU) for *in vivo* metabolic labeling of plant RNAs, which allows the isolation of neo-synthesized RNAs from intact plants. We also demonstrate that 5-EU metabolic labeling of Arabidopsis RNAs can be used for pulse-chase experiments, which allowed us to determine genome-wide Arabidopsis RNA half-lives without chemical inhibition of transcription.

## Results

### Labeling Arabidopsis RNAs with 5-Ethynyl uridine (5-EU)

First, we tested whether small molecules commonly used for *in vivo* RNA labeling exhibit toxic effects in Arabidopsis. We germinated *Arabidopsis thaliana* (Arabidopsis) wild-type Columbia-0 seed on two different concentration of the labeling substances 5-EU, BrU, or 4-SU, as well as cordycepin, an adenosine derivate commonly used to induce transcriptional arrest. Seedlings germinated on MS medium containing 4-SU or cordycepin died after germination. 5-EU or BrU had no obvious negative effect on germination or seedling development (Figure S1). We also grew seedlings on MS plates and transferred five-day old seedlings to MS plates containing 5-EU, BrU, 4-SU, or cordycepin. Seedlings transferred to 4-SU or higher concentrations of cordycepin exhibited a developmental arrest, did not develop true leaves and eventually died. In contrast, 5-EU and BrU had no obvious effect on Arabidopsis development. Higher toxicity of 4-SU compared to other labeling agents has also been observed in mammalian cell cultures (Tani et al., 2012b).

Next, we conducted a set of preliminary experiments to investigate whether feeding of the non-toxic 5-EU to Arabidopsis seedlings allows *in vivo* RNA labeling. Arabidopsis seeds were grown in liquid MS medium for five days in constant light. Then, we supplemented the growth medium with 5-EU of different concentrations. 5-EU containing RNAs were isolated from total RNA by covalent attachment of biotin to 5-EU residues followed by purification using streptavidin-coated magnetic beads. As a negative control, we isolated RNAs from seedlings that were not treated with 5-EU, but were subjected to the purification strategy for 5-EU labeled RNAs. RNAs attached to streptavidin beads were used for on-bead cDNA synthesis and subsequent RT-PCR analysis (Figure 1A).

**Figure 1:**
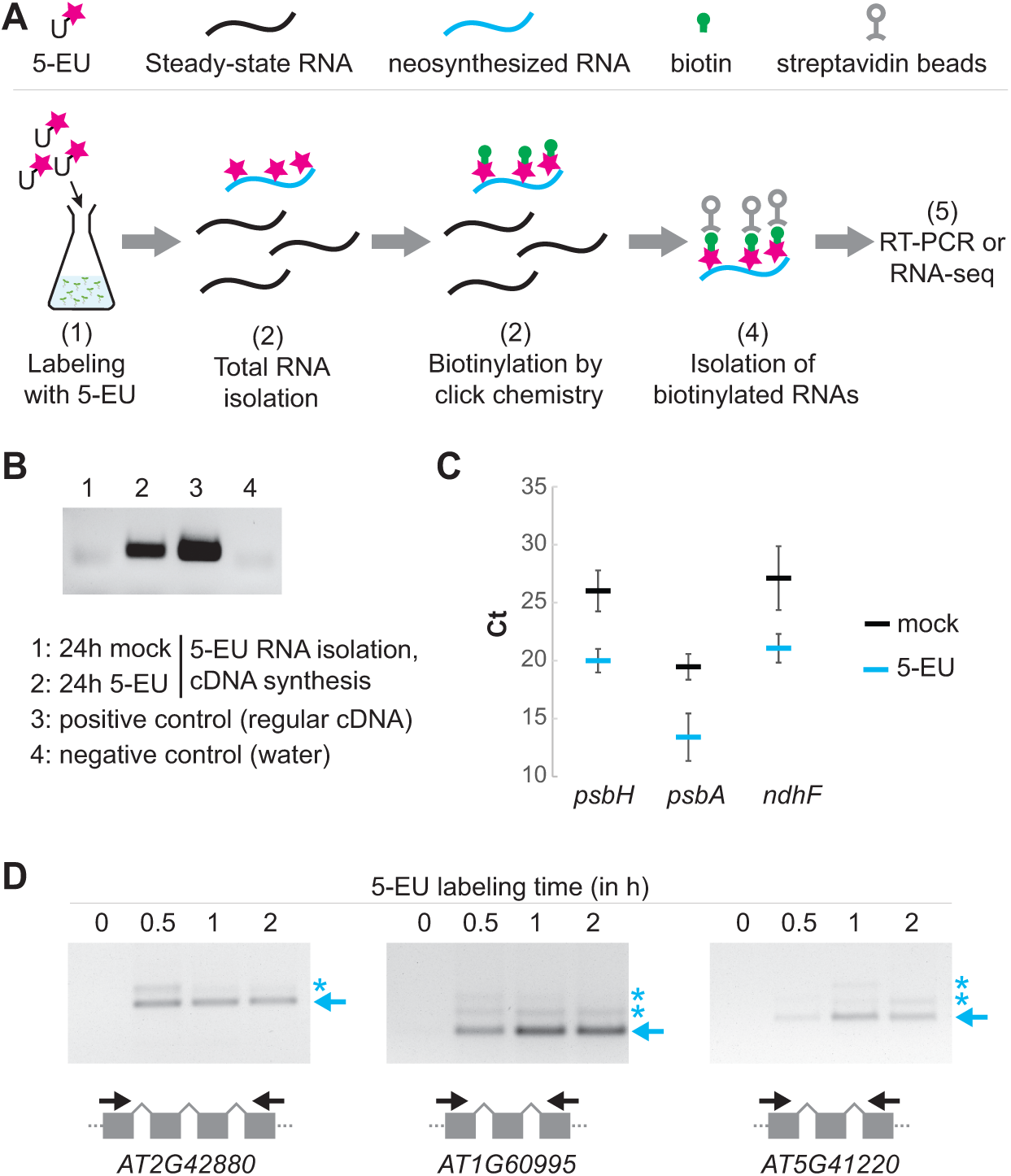
*In vivo* RNA labeling with 5-ethynyl uridine (5-EU) **(A)** Schematic presentation of the workflow for RNA labeling with 5-EU in five-day old Arabidopsis seedlings grown in liquid MS medium. **(B)** RT-PCR analysis using RNAs isolated from mock and 5-EU (200 µM) treated Arabidopsis seedlings according to the workflow shown in panel A. A conventional cDNA synthesis served as a positive control. Oligonucleotides for RT-PCR were specific for the *ACTIN2* mRNA. **(C)** RNAs isolated as in B were analyzed by qRT-PCR using oligonucleotides specific for chloroplast genes. Mean cycle threshold values are shown for the amplification of the chloroplast *psbH*, *psbA* and *ndhF* mRNA together with the corresponding standard deviations calculated based on three biological replicates. **(D)** RT-PCR analysis using RNAs from Arabidopsis seedlings treated for 0, 0.5, 1 and 2 hours with 500 µM 5-EU. Three genes were amplified with oligonucleotides flanking two or three exons. Asterisks indicate unspliced pre-mRNAs.

We found that enrichment of RNA from plants treated with 200 µM 5-EU was much higher compared to the negative control, suggesting that we achieved labeling of endogenous RNAs in *A. thaliana* seedlings with 5-EU (Figure 1B). In a separate set of replicate experiments, we also tested for enrichment of chloroplast RNAs by qRT-PCR. We found up to 70-fold enrichment of three selected mRNAs after 5-EU treatment relative to untreated controls (Figure 1C; Supplemental Table 1). Short-term pulse experiment with 500 µM 5-EU showed that 5-EU containing RNAs accumulated already 30 - 60 minutes after the addition of 5-EU to the growth medium (Figure 1D). Using oligonucleotide pairs that flank introns revealed that spliced, mature mRNAs as well as unspliced pre-mRNAs accumulated after 5-EU labeling (Figure 1D). These results suggest that 5-EU does not only permit labeling of RNAs *in vivo*, but also allows enrichment of RNA processing intermediates. Furthermore, it indicates that the plant splicing machinery processes 5-EU labeled RNAs.

### Comparison of the nascent and steady-state transcriptome

Because we observed efficient labeling of RNAs 60 minutes after addition of 5-EU to the plant growth medium (Figure 1), we chose this time point for preparation of strand-specific RNA-seq libraries. Such short-term pulses might allow the isolation of nascent pre-mRNAs and unstable RNAs, which are difficult to detect in the presence of other stable RNAs in steady-state transcriptomes. For the preparation of <u>n</u>ascent 5-<u>EU</u> labeled RNA <u>seq</u>uencing (Neu-seq) libraries, we purified 5-EU labeled RNAs from total RNAs as described above. The same total RNA sample was also used for the preparation of regular RNA-seq libraries, which we directly compared to a 60 minutes transcriptome. Because nascent pre-RNAs might not have been polyadenylated, we selectively removed rRNA and randomly primed all RNAs for RNA-seq library preparation (see Material and Methods for details). Experiments were performed in triplicates.

In order to compare the steady-state transcriptomes and Neu-seq transcriptomes, we first calculated the transcript abundances for all genes. Transcript per million (TPM) values from the steady-state and the Neu-seq transcriptome exhibited a high correlation (Figure 2A). To test whether the Neu-seq library is enriched in rare RNAs, we considered all genes with an expression value < 1 TPM as rare. When we compared Neu-seq and the regular RNA-seq libraries, we found 16,406 genes with TPM ≥ 1 in both library types (Figure 2B, Supplemental Data Set 1). The steady-state transcriptome contained only 69 RNAs (TPM > 1), which were not detected by Neu-seq (TPM < 1) (Figure 2B, Supplemental Data Set 1). Among them, we found several small nuclear RNAs (snRNAs), which are components of the spliceosome (e.g. At2g04375, At1g05057, At5g61455, At2g04455, At4g06375, At5g04085). Such RNAs are probably not produced at high rates, but exhibit high stability and are hence better detectable in steady-state transcriptomes. On the contrary, the Neu-seq library contained 4,058 RNAs (TPM > 1) that were not detectable in the regular RNA-seq library (TPM < 1) (Figure 2B, Supplemental Data Set 1). Such RNAs might be nascent or unstable RNAs, the detection of which is difficult in steady-state transcriptomes. Gene classification analysis using MapMan revealed that genes detected as expressed specifically in the Neu-seq library were enriched in e.g. non-coding RNAs including microRNAs (miRNAs) (Figure 2C) (Thimm et al., 2004). MicroRNAs are derived from longer primary RNA (pri-miRNAs). In plants, the RNAse III-like enzyme DICER-LIKE (DCL) 1 rapidly processes pri-miRNAs into mature miRNAs (Achkar et al., 2016). Pri-miRNAs are low abundant and therefore difficult to detect. In total, the expression of 30 *MIRNAs* could be detected in Neu-seq transcriptomes, but not in the steady-state transcriptome (example provided in Figure 2D, Supplemental Data Set 1), emphasizing the ability of Neu-seq to detect low-abundant RNAs such as pri-miRNAs.

**Figure 2:**
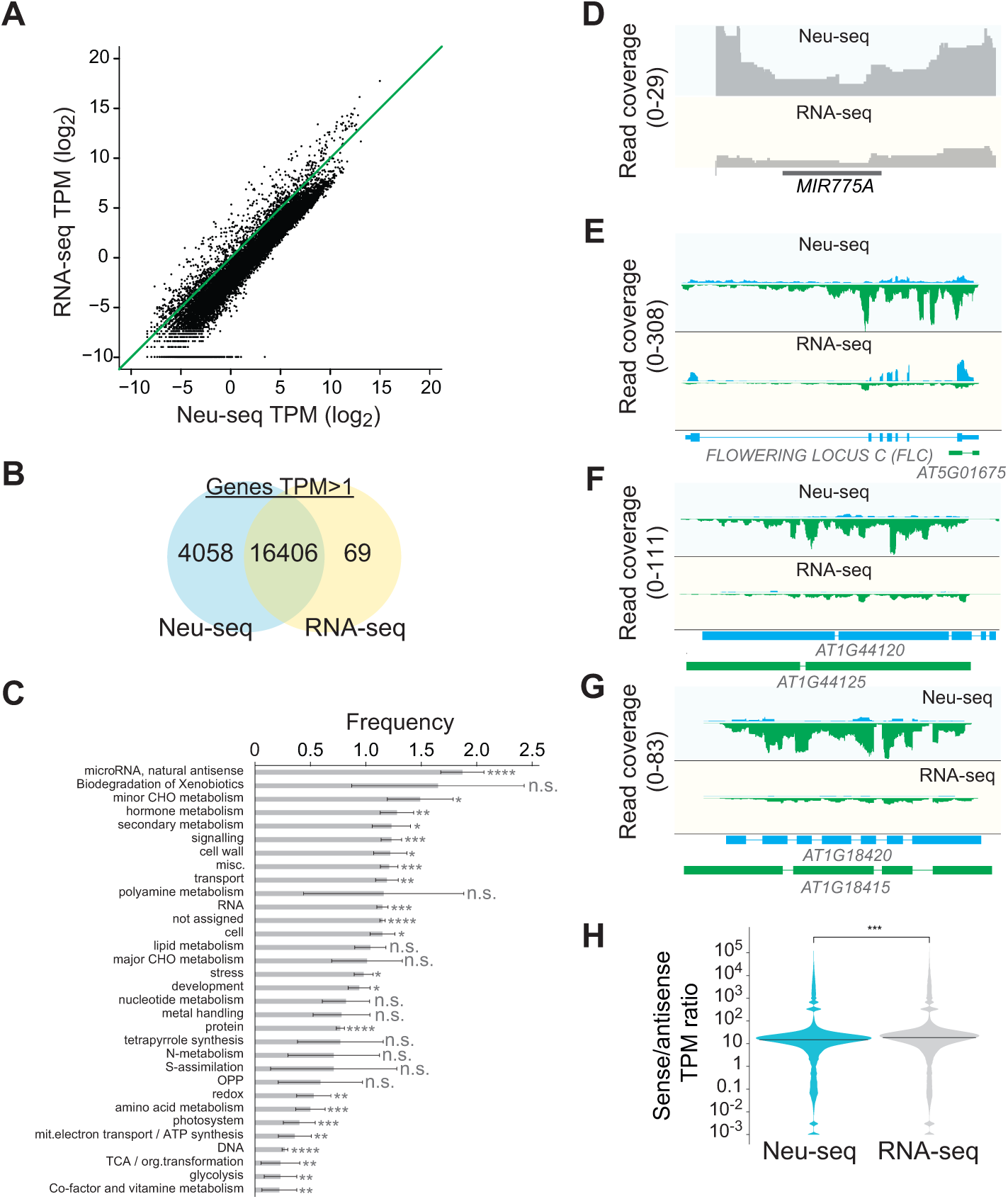
Nascent 5-EU labeled RNA sequencing (Neu-seq) detects rare RNAs including pri-miRNAs and antisense RNAs. **(A)** Comparison of transcript per million (TPM) of all genes detected by RNA-seq and Neu-seq. Arabidopsis seedlings were treated for one hour with 5-EU. The same rRNA-depleted RNAs served as material for the preparation of RNA-seq and Neu-seq libraries. **(B)** Venn diagram depicting the number of genes with TPM > 1 detected by Neu-seq and RNA-seq. **(C)** Characterization of 4058 genes for which expression was specifically detected by Neu-seq. Gene functional characterization was performed with MAPMAN. Error bars denote standard deviation. **** p < 0.0001, *** p < 0.001, ** p < 0.01, * p < 0.05, n.s. not significant **(D)** Coverage plot showing the RNA sequencing reads in Neu-seq and RNA-seq libraries mapping to the *MIR775A* gene. **(E-G)** Coverage plot showing the RNA sequencing reads in Neu-seq and RNA-seq libraries mapping to the sense and antisense strand at the *FLC* locus (E), *AT1G44120* (F), and *AT1G18420* (G). **(H)** TPM ratio of sense and anti-sense gene expression in Neu-seq and RNA-seq libraries.

Neu-seq libraries contained also more antisense RNAs than steady-state transcriptomes (Figure 2E-G, Supplemental Data Set 1). The function of antisense transcription at individual loci has been described for some Arabidopsis genes. One of the most prominent examples is *FLOWERING LOCUS C* (*FLC*). For *FLC*, transcription in antisense direction is important for the establishment of repressive chromatin marks (Whittaker and Dean, 2017). Interestingly, we observed high accumulation of antisense RNAs at the *FLC* locus in the nascent RNA transcriptome, but not in the steady-state transcriptome (Figure 2E). These results suggest that RNA production at the *FLC* locus under our experimental conditions occurs mainly in antisense direction. Similar results were obtained for other annotated antisense genes (Figure 2F, G). To test whether antisense RNAs globally accumulate in the Neu-seq libraries compared to the steady-state transcriptome, we analyzed the amounts of RNA-sequencing reads mapping in antisense orientation to annotated genes and calculated antisense TPM values for all genes. The ratio between sense/antisense TPMs was clearly shifted to sense TPMs, indicating that the majority of Arabidopsis genes is mainly transcribed in sense direction (Figure 2H). However, the ratio of sense/antisense TPMs was significantly lower in the Neu-seq library compared to the steady-state transcriptome, further indicating that Neu-seq represents antisense RNAs more comprehensively (Figure 2H). Previous Gro-seq experiments also detected substantial amounts of antisense transcripts in the Arabidopsis transcriptome (Hetzel et al., 2016). These results imply that antisense RNAs are components of the Arabidopsis transcriptome, but their detection requires more sophisticated RNA library preparation techniques such as Neu-seq or GRO-seq.

### RNA processing intermediates accumulate in the nascent transcriptome

Apart from rare RNAs we also analyzed whether Neu-seq is suitable for the detection of RNA processing intermediates such as unspliced pre-mRNAs. To investigate whether Neu-seq libraries are enriched in pre-mRNAs, we first classified reads in exonic, intronic, exon-intron junction and intergenic reads using FeatureCounts (see Material and Methods for details). We found that Neu-seq libraries contained 2.5 times more reads mapping to intron-exon boundaries (6.8% of all reads) when compared to steady-state RNA-seq libraries (2.8% of all reads; Figure 3A, B). These results suggest that Neu-seq specifically covers unspliced pre-RNAs. Reads mapping exclusively to exonic or intronic regions were not drastically changed between the two libraries (Figure 3A, B). The observation that purely intronic reads were not more abundant in Neu-seq libraries than in RNA-seq is probably due to the fact that Arabidopsis introns are relatively short. Therefore, we expect that retained introns of unspliced pre-RNAs are represented by exon-intron junction reads. The amounts of reads mapping to intergenic regions were low in Neu-seq and steady-state RNA-seq libraries (Figure 3B), which implies that the vast majority of transcribed regions in the Arabidopsis is annotated and only very few additional transcribed regions can be detected in Arabidopsis.

**Figure 3:**
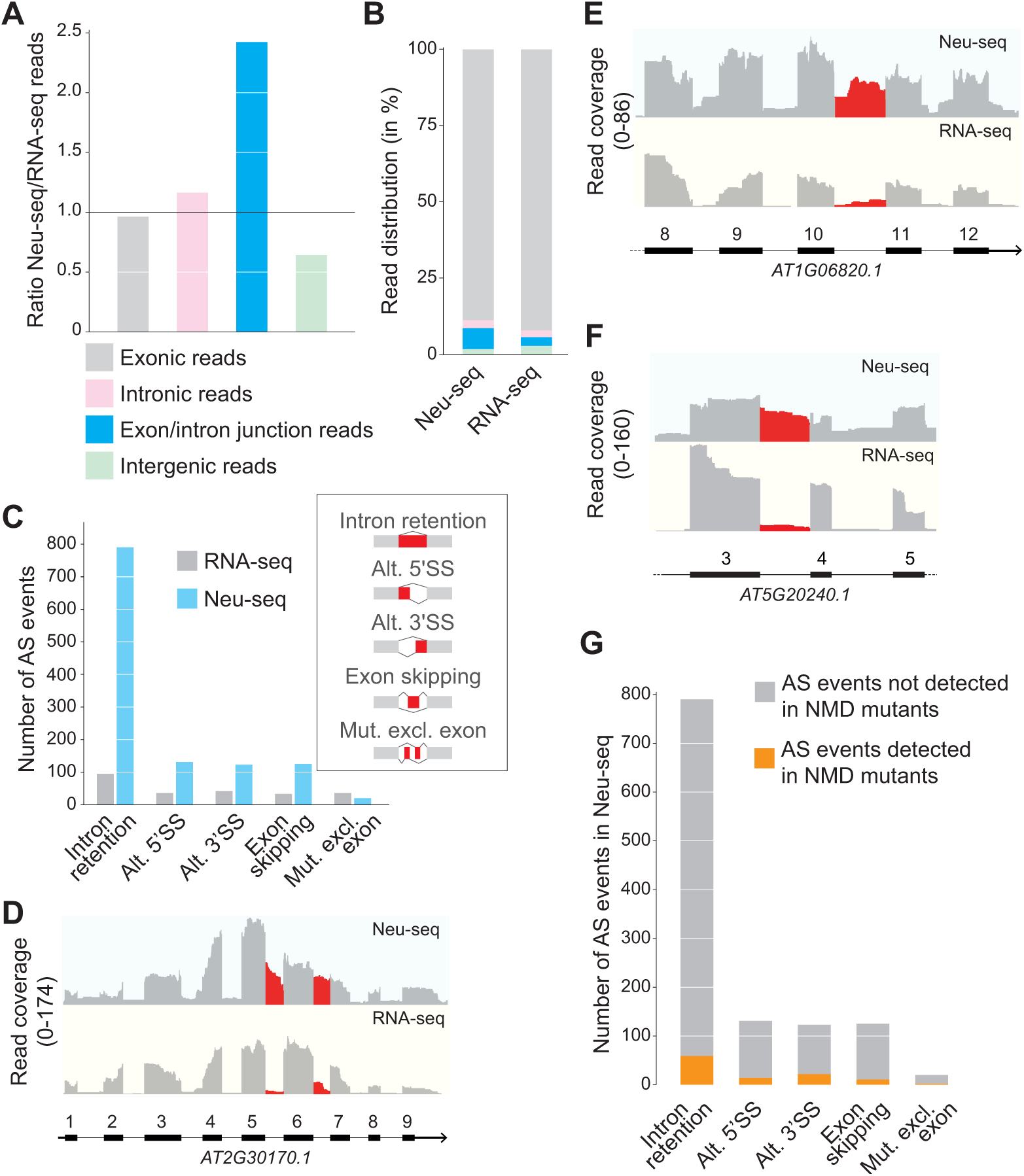
Neu-seq enriches RNA processing intermediates. **(A)** Ratio between reads in Neu-seq and RNA-seq libraries that map to exons, introns, exon/intron junctions, or intergenic regions. **(B)** Percentage of reads in Neu-seq and RNA-seq libraries that map to exons, introns, exon/intron junctions, or intergenic regions. **(C)** Detection of alternative splicing events in Neu-seq and RNA-seq libraries by rMATS. **(D-F)** Coverage plots showing the RNA sequencing reads in Neu-seq and RNA-seq libraries. Genes depicted here were identified by the rMATS algorithm as genes containing retained introns. **(G)** Overlap of Neu-seq-specific and NMD-specific alternative splicing events. Alternative splicing events detected in NMD-deficient mutants (Drechsel et al., 2013) were compared to alternative splicing events detected by Neu-seq.

Because we observed more reads mapping to exon-intron borders in Neu-seq libraries, we applied the rMATS algorithm to detect splicing isoforms that specifically accumulated in the nascent or steady-state transcriptome (Supplemental Data Set 2). The majority of the differentially spliced RNA isoforms contained retained introns, detected predominately in the Neu-seq data set (Figure 3C). Only a small number of RNAs containing retained introns were more abundant in the steady-state transcriptome (Figure 3C). Examples of introns found by the rMATS pipeline to be significantly retained in the nascent transcriptome dataset are shown in Figure 3 D-F. The rMATS pipeline allowed us to detect splicing isoforms including alternative 5’ or 3’ splice sites, mutually exclusive exons or skipped exons. We detected more RNA containing alternative 5’ or 3’ splice sites or skipped exons in the nascent transcriptome (Figure 3C). These results suggest that Neu-seq can be applied for the detection of splicing intermediates and variants.

We envisioned two distinct scenarios, which could explain the accumulation of RNAs with retained introns in the Neu-seq data set. First, such RNAs might contain introns that are spliced less efficiently compared to other introns. In that case, the splicing variants detected in the Neu-seq libraries are splicing intermediates, which will be further processed into mature mRNAs. The second alternative could be that alternative RNA isoforms in the Neu-seq dataset might be non-functional products of RNA splicing that accumulate only transiently and are subject to fast degradation. The nonsense-mediated mRNA decay (NMD) pathway removes some incorrectly spliced mRNAs and thereby regulates the expression and degradation of thousands of mRNAs in Arabidopsis (Drechsel et al., 2013; Karousis et al., 2016). In order to distinguish between the two scenarios, we compared Neu-seq data with NMD targets which were identified by transcriptome analysis of NMD mutants (Drechsel et al., 2013). We found that 7% of the genes producing RNAs with retained introns accumulated also in NMD mutants. 93% of all splicing variants containing retained introns identified by Neu-seq did not overlap with known NMD targets (Figure 3G). These observations suggest that a small subset of unspliced RNAs identified by Neu-seq is targeted for elimination by the NMD pathway, but that the majority of the RNA splicing variants detected in Neu-seq libraries might be further processed into mature mRNAs.

### Pulse-chase labeling of Arabidopsis RNAs and global analysis of RNA decay patterns

Metabolic labeling of RNA also allows stability measurements of RNA molecules by performing pulse-chase experiments. Similar to BRIC-Seq, we performed 5-<u>e</u>thynyl u<u>r</u>idine <u>I</u>P <u>c</u>hase (ERIC)-seq. To ensure high degree of labeling with 5-EU, we incubated Arabidopsis seedlings for 24 h with 200 µM 5-EU and then replaced the 5-EU containing growth media by growth media containing 20 mM unlabeled uridine. Samples were collected 0, 1, 2, 6, 12 and 24 hours after the start of the chase (Figure 4A). We isolated polyA-RNA from total RNA because we were mainly interested in the stability of mature, polyadenylated RNAs and not in RNA degradation intermediates. We spiked polyA-RNA with a 5-EU labeled *LUCIFERASE* (*LUC*) RNA (see Material and Methods for details). The *LUC* spike-in control allowed us to compare the abundance of RNAs between different samples, and to account for differences in efficiency of the click chemistry reaction, RNA purification procedures and library preparation. 5-EU labeled RNAs were purified from the polyA-RNA/*LUC* mixture as described above and used for the preparation of strand-specific RNA-seq libraries. Gene expression values were calculated and normalized to the amount of the *LUC* spike-in in the respective sample (see Material and Methods for details). Genes that exhibited an expression value of 0 TPM in any of the samples were removed. We also filtered out genes that accumulated to more than 125% after beginning of the chase. In total, 19,664 genes remained for which decay rates could be determined (Supplemental Data Set 3). To compare genes with different expression strength, we normalized all expression values relative to the expression at time point 0.

**Figure 4:**
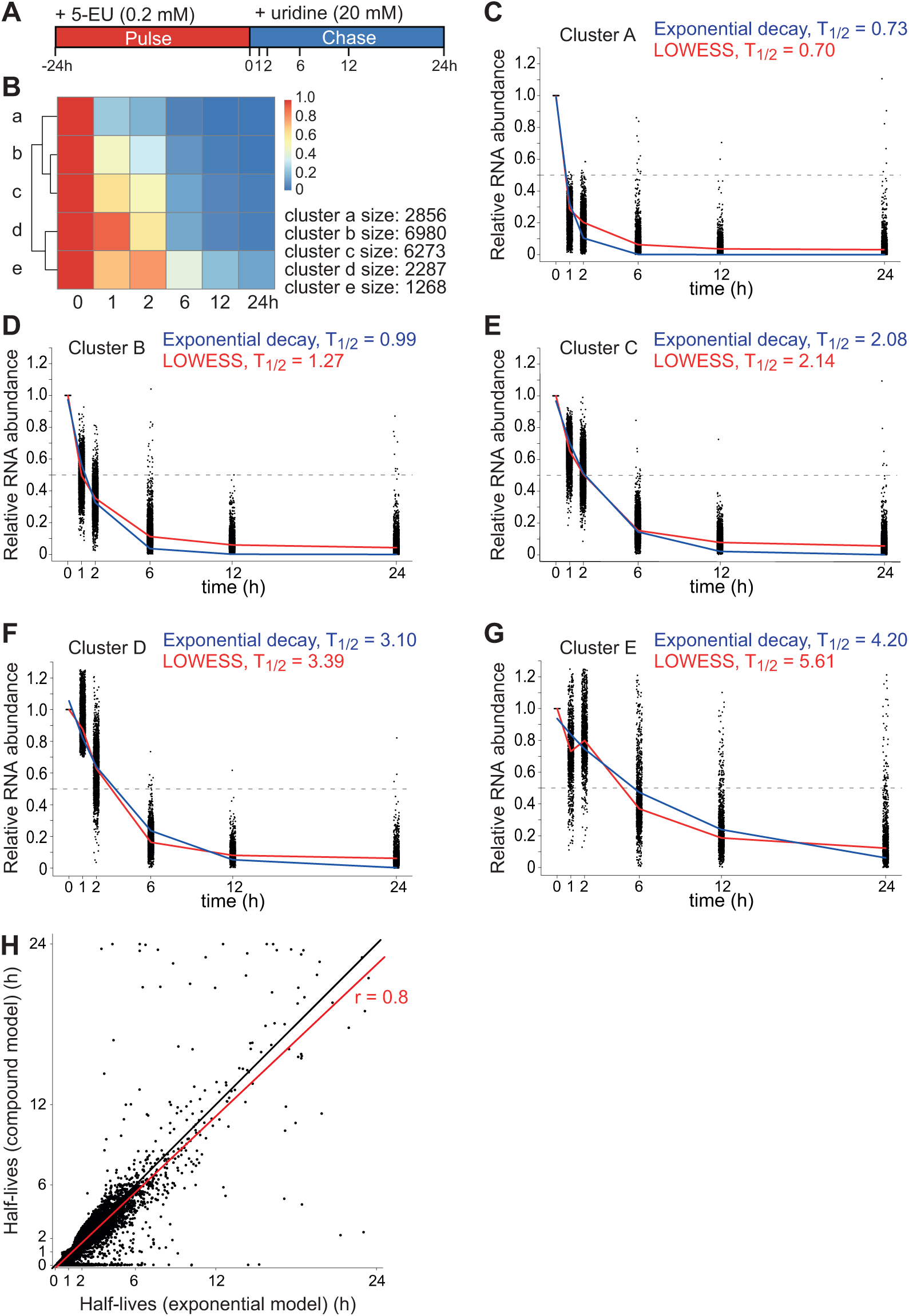
5-<u>e</u>thynyl u<u>r</u>idine <u>I</u>P <u>c</u>hase (ERIC)-seq reveals RNA half-lives in Arabidopsis seedlings. **(A)** Schematic presentation of the 5-EU pulse-chase experiment. Plants were grown for 5 days in liquid MS medium. 5-EU was added to the liquid MS and incubated for 24 hours. The liquid MS medium was replaced by liquid MS medium containing a high concentration of unlabeled uridine. Samples were taken at the indicated time points. **(B)** Hierarchical clustering of gene expression values obtained by ERIC-seq. Five forclusters exhibited distinct RNA degradation kinetics. **(C-G)** Degradation curves of all genes in cluster A-E. Relative expression values (time point 0=1) are depicted. An exponential decay model and a locally weighted polynomial regression (LOWESS) were fitted to the data points. **(H)** The half-lives of all RNAs were calculated by an exponential decay model and a compound model. The RNA half-lives estimated by the two models were correlated.

We defined five clusters of mRNAs by hierarchical clustering of gene expression values. The five different clusters exhibited distinct degradation kinetics (Figure 4B). To determine the RNA half-lives of each cluster, we applied an exponential decay model and locally weighted scatterplot smoothing (LOWESS) to calculate the half-life times *t*_1/2_ of each cluster. Genes in cluster A exhibited the fastest decay with an average half-life less than 1 h (*t*_1/2_ = 0.7) (Figure 4C). Cluster B to E contained more stable RNAs, featuring average half-lives of 1.0, 2.1, 3.1 or 4.2 hours, respectively (Figure 4D-G). The exponential decay model and LOWESS in cluster A to D yielded similar half-lives, suggesting that RNAs in this clusters decayed exponentially (Figure 4C-F). The exponential decay model and LOWESS resulted in different average half-lives in cluster E (4.2 h vs. 5.6 h). This might be caused by the unexpected abundance pattern of RNAs in this cluster: RNAs in cluster E decayed after 1 h, but re-accumulated 2 h after the chase, resulting in an internal peak (Figure 4G). This unexpected behavior might be explained by re-incorporation of 5-EU nucleotides, which are generated after degradation of 5-EU labeled RNAs. Although we chased with a 100 times higher concentration of the unlabeled nucleoside uridine, we cannot be sure that these uridine nucleosides are quickly converted into unlabeled uridine nucleotides, which are then used for transcription. High levels of re-incorporation of 5-EU nucleotides might occur especially in RNAs produced from genes with high transcriptional rate. Importantly, decay rates for genes which fall in cluster E (total number 1233) should be interpreted carefully.

We also calculated the RNA half-lives from all genes individually. We applied two different models for the calculation of half-lives, an exponential decay model and a compound model. The compound model fits the raw data by a convolution of Voigt curve and exponential decay curve (Shippony and Read, 1993). Half-life values determined by exponential and compound model fitting showed high correlation (r=0.8), but RNA half-lives determined by the compound model were shifted to shorter half-lives (Figure 4H). These results reveal that both models describe the decay rates of RNAs in a similar manner, but that the compound model predicts shorter RNA half-life values for some genes. We provide all estimated half-lives determined by compound and exponential models in Supplemental Data Set 4. In general, RNAs exhibited a wide range of predicted half-lives from minutes to a few hours (Supplemental Data Set 4). Because we found many RNAs with half-lives less than one hour, we suggest a higher timescale resolution within the first hours of the chase for future experiments.

Given that chloroplast RNAs were efficiently labeled by 5-EU, we tested whether we can detect decay by performing a separate set of experiments on samples chased with non-labelled uridine for 0, 2 and 6 hours. A synthetic, EU-labeled transcript named *aadA* served as a spike-in after RNA extraction for all samples and the data were normalized according to the *aadA* signal. We chose three mRNAs for our RNA decay analysis by qRT-PCR, which are all described to be stabilized in a protein-dependent, i.e. potentially regulated manner (Lezhneva and Meurer, 2004; Johnson et al., 2010; Kupsch et al., 2012). All three mRNAs show decreasing signal over time (Figure S2). This analysis demonstrates that RNA turnover measurements based on metabolic labeling are possible in chloroplasts.

### Comparison of mRNA half-lives determined by ERIC-seq and transcriptional arrest experiments

We compared RNA half-lives determined by ERIC-seq with previously published half-lives for Arabidopsis mRNAs determined by transcriptional arrest experiments. For this comparison, we obtained mRNA half-lives calculated in experiments using actinomycin D (Narsai et al., 2007) and cordycepin (Sorenson et al., 2018). The overall Pearson correlation of the half-lives determined by ERIC-seq and transcriptional arrest experiments were 0.31 and 0.19 (Spearman correlation 0.28 and 0.28, respectively), which indicates low correlation between the different methodologies (Figure 5A). Also the half-lives determined by Narsai et al. and Sorenson et al. exhibited only a low correlation with each other (Pearson correlation 0.33, Figure S3). These results reveal that half-lives determined by all three approaches are largely different.

**Figure 5:**
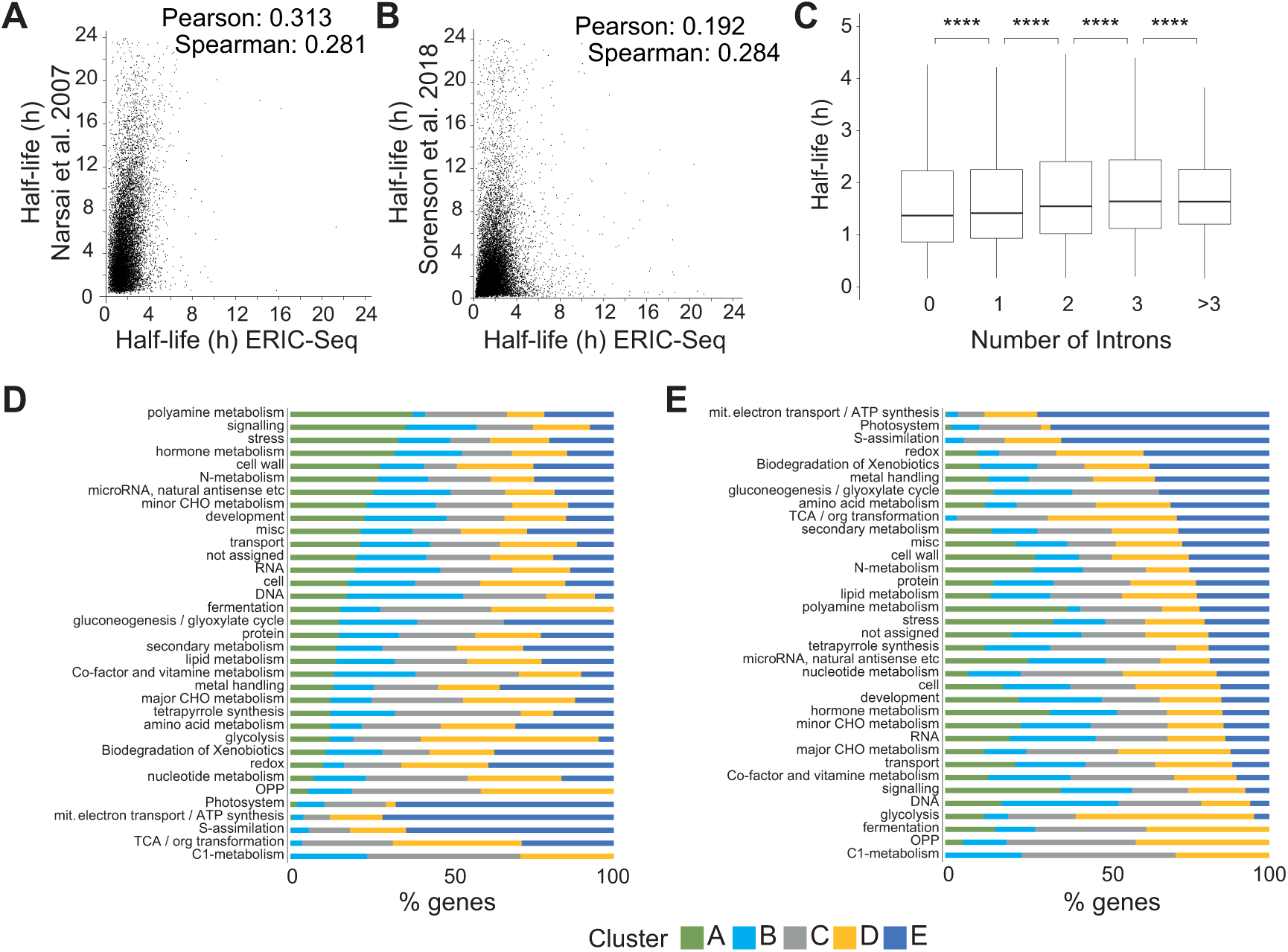
Comparison of inhibitor-free and transcriptional inhibitor-based estimations of RNA half-lives. **(A,B)** Comparison of RNA half-lives estimated by inhibitor-free (ERIC-seq) and inhibitor-based approaches. Narsai et al. applied the transcriptional inhibitor actinomycin D to Arabidopsis cell cultures for calculation of RNA half-lives (A). Sorenson et al. applied the transcriptional inhibitor cordycepin to Arabidopsis seedlings for the calculation of RNA half-lives (B). **(C)** Box plots depicting the half-lives (determined by ERIC-seq) of RNAs derived from genes with 0, 1, 2, 3 or more than 3 introns. Wilcoxon-Mann-Whitney test, ****p<0.0001 **(D-E)** Functional gene characterization by MAPMAN of genes which fall in cluster A, B, C, D or E. Functional gene categories were sorted by their abundance in cluster A, which exhibits the shortest half-life (depicted in panel D), or by their abundance in cluster E, which exhibits the longest half-life depicted in panel E.

Interestingly, RNA half-lives determined by ERIC-seq are much shorter than the ones determined after treatment with cordycepin or actinomycin D (Figure 5A, B). However, general trends deduced from previous studies were confirmed using ERIC-seq: Genes with no or few introns produced RNAs with shorter half-lives (Figure 5C). Previous studies also found that RNA stability correlates with distinct biological functions (Narsai et al., 2007, Sorenson et al., 2018). To test this, we performed a gene classification analysis using MAPMAN on the five different clusters described before (Thimm et al., 2004). Genes belonging to cluster A contained RNAs with the shortest half-lives and were enriched in non-coding RNAs, stress-related genes, genes involved in hormonal pathways and the metabolism of polyamines (Figure 5D). On the contrary, genes from cluster E, the RNAs of which exhibit the longest half-lives, were specifically enriched in genes responsible for mitochondrial and plastid functions as well as primary metabolism (such as the TCA cycle) (Figure 5E).

## Discussion

*In vivo* RNA labeling is largely used in the yeast and metazoan RNA research community, but is not well established in the plant field. We used 5-EU, a uridine homolog, for *in vivo* RNA labeling in Arabidopsis seedlings. Unlike 4-SU, we did not observe toxic effects of 5-EU treatment during germination or seedling development. In addition, our results show that 5-EU is taken up by the plant, it incorporates into RNAs generated from nuclear and chloroplast genes, and it does not interfere with RNA processing steps such as splicing. 5-EU is effectively coupled to biotin using click chemistry, which allows efficient purification of labeled RNA molecules. We found that 5-EU labeling of RNAs *in vivo* combined with RNA-seq is a powerful tool to gain important insights into the dynamics of the Arabidopsis transcriptome.

### Neu-seq: from identification of rare and unstable RNAs to measurements of transcriptional activity

We analyzed an Arabidopsis 1h-transcriptome with Neu-seq and compared it to a regular steady-state transcriptome determined by standard RNA-seq. Neu-seq detected more low abundant RNAs, including pri-miRNAs and antisense transcripts, than the standard RNA-seq experiment. Our Neu-seq libraries did not contain many more sequencing reads emerging from intergenic regions, most likely due to the high-quality annotation of the Arabidopsis genome. However, Neu-seq could still be useful for the detection of unknown transcribed regions that might be only visible in certain Arabidopsis mutants defective in RNA turnover and in other plants with much less well-annotated genomes. In general, Neu-seq provides a simple alternative to GRO-Seq and related techniques for the identification of nascent transcripts and thus can serve as a proxy for determining transcriptional activity.

Another promising application of Neu-Seq is the possibility to assess chloroplast transcription. Chloroplast transcription depends on nuclear and chloroplast-encoded RNA polymerases and is regulated in response to various cellular and external signals, often via changes in the chloroplast redox state (Pfannschmidt et al., 1999; Liere et al., 2011). Chloroplast transcriptional activity was so far assayed by run-on techniques that depend on broken chloroplasts. This disruptive treatment precludes an analysis of the impact of changing external conditions on transcription (Legen et al., 2002). The finding that chloroplast RNAs can be labeled *in vivo* with externally supplied 5-EU opens up options to determine transcriptional activity under diverse growth conditions. How 5-EU traverses the double-membraned chloroplast envelope is unknown at present. It is, however, clear that there has to be import of pyrimidines into chloroplasts, since the final steps of pyrimidine *de novo* synthesis occur in the cytosol (Witz et al., 2012). Salvage enzymes like uracil phosphoribosyl transferase as well as plastidic uridine kinases and pyrimidine degradation enzymes are found in chloroplasts and are essential for photosynthesis and plant development (Mainguet et al., 2009; Zrenner et al., 2009; Chen and Thelen, 2011; Cornelius et al., 2011). The proton antiporter PLUTO is currently the only known transport protein in *Arabidopsis* mediating the import of uracil into plastids for salvage reactions and degradation (Witz et al., 2012). Likely, 5-EU is a substrate for this transporter as well, which would explain the success of 5-EU based labeling of chloroplast transcripts

### Neu-seq for RNA processing studies

The nascent 1h-transcriptome contained 2.5 times more reads mapping to intron/exon boarders compared to steady-state transcriptomes, indicating an enrichment of unspliced pre-mRNAs in the 1h-transcriptome. mRNA isoforms specific for the nascent 1h-transcriptome showed a small overlap with known NMD targets. This suggests that the majority of mRNA isoforms specific for the 1h nascent transcriptome will be further processed into mature mRNAs. Most of the splicing isoforms detected in the nascent 1h-transcriptome still contained introns, which is in line with idea that these introns will be further spliced. We also detected more alternative 5’ and 3’ splicing events in the 1h-transcriptome than in the steady-state transcriptome. An attractive hypothesis for the emergence of alternative 5’ and 3’ isoforms is recursive splicing (Georgomanolis et al., 2016). This non-canonical splicing reaction removes long introns in a stepwise manner. The stepwise processing steps generate low-abundant splicing intermediates, which contain alternative 5’ or 3’ splice sites. The extend of recursive splicing in plants is not known, but the analysis of nascent 5-EU labeled transcriptomes in conjunction with specialized bioinformatic approaches for detecting recursive splicing will likely solve this question in the future (Pulyakhina et al., 2015). In general, Neu-seq will be a major improvement for future studies on plant RNA processing. A more detailed analysis of RNA processing kinetics will require shorter and multiple 5-EU labeling periods (Schwarzl et al., 2015). 5-EU labeling experiments can be extended for the analysis of RNA splicing in RNA processing mutants, which might shed light on the exact functions of specific proteins in regulating RNA processing (Tani et al., 2012a; Maekawa et al., 2015).

### ERIC-seq for genome-wide measurements of RNA half-lives

Discrepancies between RNA half-lives determined by different methods are very common. Possible explanations are bias introduced by each different technique for determining RNA half-lives. All methods have in common that certain molecules enter the cell to execute a specific function. The rate with which a small molecule is taken up, and the lag phase until the small molecule reaches sufficient amounts to e.g. block transcription, is different depending on the molecule. Thus, depending on the method applied and the cell line used, a wide range of different half-lives is observed. In mammals, the same mRNA exhibited half-lives from more than 10 hours in early experiments compared to only 10 minutes in later studies (Yang et al., 2003; Rabani et al., 2011a). We also observed that RNA half-lives determined by ERIC-seq dramatically differ from previous studies which applied cordycepin or actinomycin to block transcription (Narsai et al., 2007; Sorenson et al., 2018). Our results suggest that plant RNA half-lives determined by metabolic labeling experiments are in general shorter than previously reported RNA half-lives. One possible explanation for the discrepancies between transcriptional arrest and metabolic labeling experiments (such as ERIC-seq) is the crosstalk between transcription and RNA degradation, which stabilizes RNAs after blocking transcription (Sun et al., 2013; Braun and Young, 2014). Another possible explanation for the observed differences is that cordycepin and 4-SU exhibit strong toxic effects in Arabidopsis. The application of 5-EU for RNA stability studies might also introduce certain bias. For instance, we cannot exclude that 5-EU containing RNAs are less stable than native RNAs *in vivo*. But the fact that 5-EU, unlike 4-SU or cordycepin, does not exhibit toxic effects on Arabidopsis makes this explanation less likely. Apart from the methodology, also differences in plant growth and treatment conditions among the different studies might influence RNA stabilities. Nevertheless, general conclusions based on experiments with transcriptional inhibitors still hold true for the results we obtained by ERIC-seq. For instance, mRNAs with long half-lives often encode proteins involved in photosynthesis, electron transport or other basic metabolism (Narsai et al., 2007; Sorenson et al., 2018). mRNAs with short half-lives contain fewer introns than long-lived RNAs and often encode proteins involved in signal transduction, stress defense or hormone signaling (Narsai et al., 2007; Sorenson et al., 2018). Given these similar patterns, we would argue that RNA half-lives measured by transcriptional arrest based experiments yield valuable information, but may greatly overestimate RNA stability. What the true half-live of an Arabidopsis mRNA is might be accessible by a future combination of genome-wide techniques calibrated by single-molecule analyses like the recently established single RNA imaging and turnover assays (Horvathova et al., 2017)

### Summary and final thoughts

In this article, we describe the versatility of 5-EU for *in vivo* RNA labeling in Arabidopsis seedlings and its application in mRNA detection, processing and stability analyses. One advantage of our method is that the purification of 5-EU labeled RNAs is performed using a commercially available kit. Hence, many researchers can adopt this method without the need for protocol optimization. This technique allows to study RNA production and stability under different conditions such as temperature stress or hormone treatments. Moreover, 5-EU labeling experiments can be extended for the analysis of RNA splicing and processing mutants and help to understand the exact function of a specific protein in regulating RNA metabolism (Tani et al., 2012a; Maekawa et al., 2015). In the future, Neu-seq and ERIC-seq could be adapted for RNAs, which require specialized library preparation protocols such as miRNAs or circular RNAs (Tani et al., 2015; Marzi et al., 2016). ERIC-seq combined with methods for accurate mRNA isoform detection, including 3’end sequencing, will allow to determine RNA stability of specific isoforms. We are looking forward to other researchers using 5-EU *in vivo* labeling of RNAs to address yet-unanswered scientific questions in the field of plant RNA metabolism.

## Material and Methods

### Plant material, growth conditions and 5-EU treatment

*Arabidopsis thaliana* wildtype seed (Col-0) was surface-sterilized using 70% ethanol. For toxicity assays, we grew Arabidopsis on half-strength MS medium containing the indicated amount of 5-EU, BrU, 4-SU and cordycepin. Alternatively, we grew Arabidopsis on half-strength MS medium for five days and then transferred the seedlings to half-strength MS medium containing 5-EU, BrU, 4SU and cordycepin. Plants were grown under long-day conditions (16h light/8h darkness) for 14 days.

For 5-EU labeling experiments, 25 µg of seeds were added to 50 ml half-strength MS in an 125 ml Erlenmeyer flasks and stratified for two days at 4°C in the dark. Plants were grown in constant light with constant shaking on a platform shaker (130 rpm). Seedlings grew for five days, and were then separated into triplicates. For Neu-seq experiments, seedlings were treated with 500 µM 5-EU for 1h and snap-frozen in liquid nitrogen. For the ERIC-seq and qRT-PCR experiments, seedlings were treated with 200 µM 5-EU for 24 h. After the 5-EU treatment, seedlings were washed three times using half-strength MS medium, and then placed in half-strength MS medium containing 20 mM uridine. Aliquots of the seedlings were harvested after 0, 1, 2, 6, 12 and 24 h and snap-frozen in liquid nitrogen.

The protocol was adjusted for tests of chloroplast RNA labeling and turn-over. First, 30-35 mg of Arabidopsis Col-0 seeds were used for surface sterilization with 70% ethanol. These were stratified in 40 ml half-strength MS medium in a culture flask as described above. For the +EU experiments, two flasks with five days old seedlings were pooled for one replicate experiment. For the –EU control experiments only one flask was used per replicate. Aliquots of 5-EU seedlings were harvested after 0, 2, and 6 h. Control reactions without label were performed for the 0 h time point.

### RNA extraction

RNA was extracted using the RNeasy Plant Mini Kit (Qiagen) for total RNA analysis and with the Direct-zol RNA Miniprep Kit (Zymo Research) for chloroplast RNA analysis according to the manufacturer’s protocols. Remaining genomic DNA in the RNA samples was removed by DNAse treatment (Thermo Fisher Scientific) followed by clean-up with the RNA Clean and Concentrator Kit (Zymo Research) according to the manufacturer’s protocol. For chloroplast RNA analysis, TURBO DNase (Thermo Fisher Scientific) treatment was performed before the cDNA synthesis and DNase was heat inactivated (see qRT-PCR section).

### Isolation of 5-EU labeled nascent RNA and Illumina library preparation

For Neu-seq libraries, rRNA were removed from 10 µg of DNAse-treated RNA using the RiboZero according to the manufacturer’s protocol (Illumina). 30 ng rRNA-depleted RNA samples were used for steady-state transcriptome library preparation and the remaining RNA was subjected to isolation of 5-EU labeled RNA. We used the Click-it® Nascent RNA Capture Kit (Life Sciences) for the attachment of biotin to the 5-EU containing RNA according to the manufacturer’s recommendations. In brief, biotin molecules were covalently attached to 5-EU. Streptavidin-coated magnet beads specifically bind biotinylated RNAs and un-labeled RNAs were washed away. 5-EU labeled RNAs bound to beads and rRNA-depleted total RNAs were used as starting material for Total RNA Illumina® ScriptSeq Plant Leaf Kit (Illumina) according to manufacturer’s recommendations.

For pulse-chase experiments, polyA-RNAs were isolated from 5 µg of DNAse I treated RNAs using the NEB Next® Poly(A) mRNA Magnetic Isolation Module according to the manufacturer’s protocol except that RNAs were eluted from the beads in RNAse-free water. Each sample was spiked with 10 pg 5-EU-labeled *LUCIFERASE* RNA (see below) and subjected to library preparation as described above.

The protocol for isolation of chloroplast RNAs followed the above description closely, with the following deviations: For the qRT-PCR experiments, 2.5 µg total RNA was used for the attachment of biotin. Each sample was spiked with 0.1 ng 5-EU labeled *aadA* RNA (see below). The immobilized beads were additionally incubated for 5 min on a rotator in Wash Buffer 1 and subsequently in Wash Buffer 2, before the standard washing steps of the Click-it® protocol. Furthermore, low nucleic acid binding tubes were used for all incubations. For two of the three replicates, instead of the Dynabeads® MyOne™ Streptavidin T1 magnetic beads of the Click-it® kit, Dynabeads® MyOne™ Streptavidin C1 magnetic beads (Thermo Scientific) were used.

### qRT-PCR analysis

5-EU-labeled (+EU) and non-labeled RNAs (-EU) bound to the beads were resuspended in RNAse-free water and treated with TURBO DNase (Thermo Scientific). The DNAse was heat inactivated and reverse transcription was performed with random hexamer primers using the RevertAid First Strand cDNA Synthesis Kit (Thermo Scientific) according to the manufacturer’s protocol. The cDNA was diluted 1:10 for control and 1:100 for time course experiments. qRT-PCR was performed with a oligonucleotide concentration of 300 nM using the Luna Universal qPCR Master Mix (NEB). The results were analyzed using the Applied Biosystems 7500 Fast Real-Time PCR Software v2.3 (Thermo Fisher Scientific). All oligonucleotides used in this study are listed in Supplementary Table 2 and were described before (de Longevialle et al., 2008; de Francisco Amorim et al., 2018).

### In vitro transcription of 5-EU labeled spike-in RNAs

For the generation of 5-EU labeled *LUC* RNA, we used a control plasmid provided in the TnT® Quick Coupled Transcription/Translation System (Promega). The plasmid was linearized using the restriction enzyme *Sac*I and purified using the DNA Clean and Concentrator Kit (Zymo Research). We used the TranscriptAid T7 High Yield Transcription Kit (Thermo Scientific) according to the manufacturer’s protocol for the “Synthesis of Non-Radioactively Labeled RNA Probes”. 5-Ethynyl-UTP was purchased from Jena Bioscience. A fraction of the reaction was analyzed using a standard agarose gel. The RNA was purified with the RNA Clean and Concentrator Kit (Zymo Research).

As a spike-in control for the chloroplast RNA turnover analysis, an *aadA* PCR product was amplified using a reverse oligonucleotide including the T7 promoter and purified using the GeneJET PCR Purification Kit (Thermo Scientific). The 5-EU labeled *aadA* spike-in was transcribed *in vitro* from the PCR product using the T7 RNA Polymerase (Thermo Scientific). Labeling and purification were carried out as for the *LUC* RNA.

### Illumina read processing of Neu-seq libraries

Single end libraries of Neu-seq and steady-state transcriptomes (three replicates each) were sequenced on an Illumina Hiseq3000. We gained a total of 381,127,501 reads. Adapters were removed and reads were trimmed of low-quality bases using Trimmomatic version 0.36 with the following parameters: ILLUMINACLIP:TruSeq3-SE.fa:2:30:10 LEADING:3 TRAILING:3 SLIDINGWINDOW:4:15 MINLEN:35 (Bolger et al., 2014). Ribosomal and tRNA reads were filtered out using Bowtie2 version 2.2.9 with the following parameters --fr--un-gz (Langmead and Salzberg, 2012). tRNA and rRNA sequences were retrieved from a tRNA database collection (Chan & Lowe, 2009) (http://gtrnadb2009.ucsc.edu/download.html), from NCBI geneBank (ID: X52320.1), and TAIR (AT2G01010.1 – 18SrRNA, AT2G01020 – 5.8sRNA, ATCG00920 – 16SrRNA, ATCG00950 – 23SrRNA, ATCG00960 – 4.5SrRNA, ATCG01160.1 – 5SrRNA, ATMG00020.1 – 26SrRNA). 372,364,707 reads survived filtering and trimming. Filtered reads were mapped to the *Arabidopsis thaliana* Ensembl genome reference (release 36) using Tophat2 version 2.1.1 with the following parameters: -p 10 -i 10 -I 1000 --library-type fr-secondstrand with an average 83 % mapping rate (Kim et al., 2013; Kersey et al., 2016). All sequencing data were publicly deposited (accession number GSE118462, Gene Expression Omnibus)

### Analysis of Neu-RNAseq libraries

For analyzing steady-state RNA-seq and Neu-seq libraries, we developed a pipeline consisting of library processing, mapping to the reference genome and analyzing read count composition of exonic and intronic features. Mapped reads were collected with featureCounts from the RSubread package version 1.5.2 with the following parameters: -T 6 -F SAF -J –O and gene coordinates or exon and intron coordinates as features (Liao et al., 2013). TPM and log_2_ TPM values were calculated from collected feature read counts. We added 0.001 to every value to avoid 0 read counts. Log_2_ TPM values of features coming from the Neu-seq and steady-state transcriptome libraries were plotted with ggplot version 2.2.1 (Wickham, 2009). Spanning read composition analysis was performed using featureCounts’ “detailed read assignment” result files. In these files, reads were collected and categorized to exonic, intronic and exonic-intronic categories based on their position among features. Average values from triplicate measurements from Neu-seq and steady-state transcriptome libraries were compared using a Fisher’s exact one-tailed test in R (R Core Team, 2017).

### Analysis of alternative splicing

For analyzing alternative splicing events, mapped libraries (after removal of tRNA and rRNA contaminations) were analyzed using rMATS version 3.2.5 applying the following parameters: -t single -len 150 (Shen et al., 2014). Events of exon skipping, mutually exclusive exon usage, alternative 5’ or 3’ splice sites and intron retentions were extracted based on p-value < 0.05 and FDR 0.05. Detected alternative splicing events are listed in Supplementary data set 2.

### Illumina read processing of ERIC-seq libraries

Sequencing libraries of 5-EU-labeled RNAs 0, 1, 2, 6, 12 and 24 h after the chase (three replicates each) were sequenced on an Illumina Hiseq3000. A total of 395,194,241 reads were sequenced. Adapter removal, trimming of low-quality bases, and tRNA and rRNA filtering and mapping was performed as described above. In total, 339,411,658 reads remained for further analysis. All sequencing data were publicly deposited (accession number GSE118462, Gene Expression Omnibus)

### Analysis of ERIC-seq libraries

Mapped reads were collected with featureCounts from the RSubread package version 1.5.2 with the following parameters: -T 6 -F SAF -J –O (Liao et al., 2013). Differential expression analysis was performed using DESeq2 version 1.16.1 with substituting size factors of estimated size factors from the luciferase gene read counts (Love et al., 2014). All genes were removed from downstream analysis that showed a zero-expression value in any of the replicates. Shrunken log_2_ fold changes of the expression values were extracted from the DEseq2 object. Additional normalization of shrunken log_2_ values was executed. Time point value changes were calculated relative to time point 0 (T0 = 1). If the expression of a gene exceeded 1.25 in any of the later time points, the gene was removed from further analysis. Normalized and filtered values were used for visualizing using pheatmap version 1.0.8 and model fitting (Kolde, 2012).

Based on the categorization of five separate clusters of the heatmap analysis, locally-weighted polynomial regression (LOWESS) and exponential model decay fitting were performed on the five clusters in R to compare overall average half-life values using the following formulas: lowess(x=X, y=Y, f=0.2) and nls(Y ∼ a*exp(-k * X), time_course_object, start=c(a=1, k=1)). The calculation of individual half-life values of all genes was performed using an exponential decay model or a composite model with an exponential background and Voigt peak model. All calculations were performed using the python lmfit package version 0.9.7 (Newville, 2016).

### Visualization of RNA-seq results

For visualization snapshots of the RNA-seq data we used the Integrative Genomics Viewer version 2.3.80 (Robinson et al., 2011; Thorvaldsdóttir et al., 2013).

## Supporting information

Supplemental Data Set 1

Supplemental Data Set 2

Supplemental Data Set 3

Supplemental Data Set 4

## Acknowledgements

Technical support by Irina Passow is gratefully acknowledged. This work was supported by the DFG (TRR175-A02 to CSL, LA2633-4/1 to S.L.), and by the Max Planck Society Chemical Genomics Centre II through its supporting companies AstraZeneca, Bayer CropScience, Bayer Healthcare, Boehringer-Ingelheim, and Merck (to S.L.). We thank all members of the lab for critical reading of the manuscript.

## Author contributions

E.X.S., P.R., M.-K. L., U.G., C.S.-L. and S.L. designed the research. P.R., M.-K. L., M.O. and M.d.F.A. performed research. E.X.S., P.R., M.-K. L., U.G., C.S.-L. and S.L. analyzed data. E.X.S., P.R., M.-K. L., C.S.-L. and S.L. wrote the paper with contributions from all authors.

---

**Figure S1: Effects of bromouridine (BrU), 5-Ethynyl Uridine (5-EU), 4-thiouridine (4-SU), and cordycepin (cor) on plant development**

**(A)** Arabidopsis plants grew on plant half-strength MS medium containing the indicated amounts of BrU, 5-EU, 4-SU, and cordycepin for 14 days under long-day conditions.

**(B)** Arabidopsis plants grew for five days on plant half-strength MS medium. Seedling were transferred to half-strength MS medium containing the indicated amounts of BrU, 5-EU, 4-SU, and cordycepin. Seedlings developed for another 14 days under long-day conditions.

**Figure S2: Metabolic labeling for RNA stability measurements of chloroplast RNAs**

Decay of selected chloroplast mRNAs measured after 24 hours of 5-EU labeling and subsequent chase with non-labeled uridine. Relative expression values (time point 0=1) are shown. For the analysis of *ndhF* RNA decay, three biological replicates were performed, for *psaA* and *rbcL* two. Graphs represent exponential trend lines with curve equations and R square values displayed in the graph areas.

**Figure S3: Comparison of RNA half-lives estimated by inhibitor-based approaches.**

Comparison of RNA half-lives determined by Narsai et al. and Sorenson et al.. Narsai et al. applied the transcriptional inhibitor actinomycin D to Arabidopsis cell cultures. Sorenson et al. applied the transcriptional inhibitor cordycepin to Arabidopsis seedlings.

**Supplemental Table 1: qRT-PCR analysis of chloroplast mRNAs after 5-EU labeling**

**Supplemental Table 2: List of all oligonucleotides**

**Supplemental Data Sets 1-4**

