## Supplementary material for "Metabolic labeling of RNAs uncovers hidden features and dynamics of the *Arabidopsis thaliana* transcriptome": Figure S3

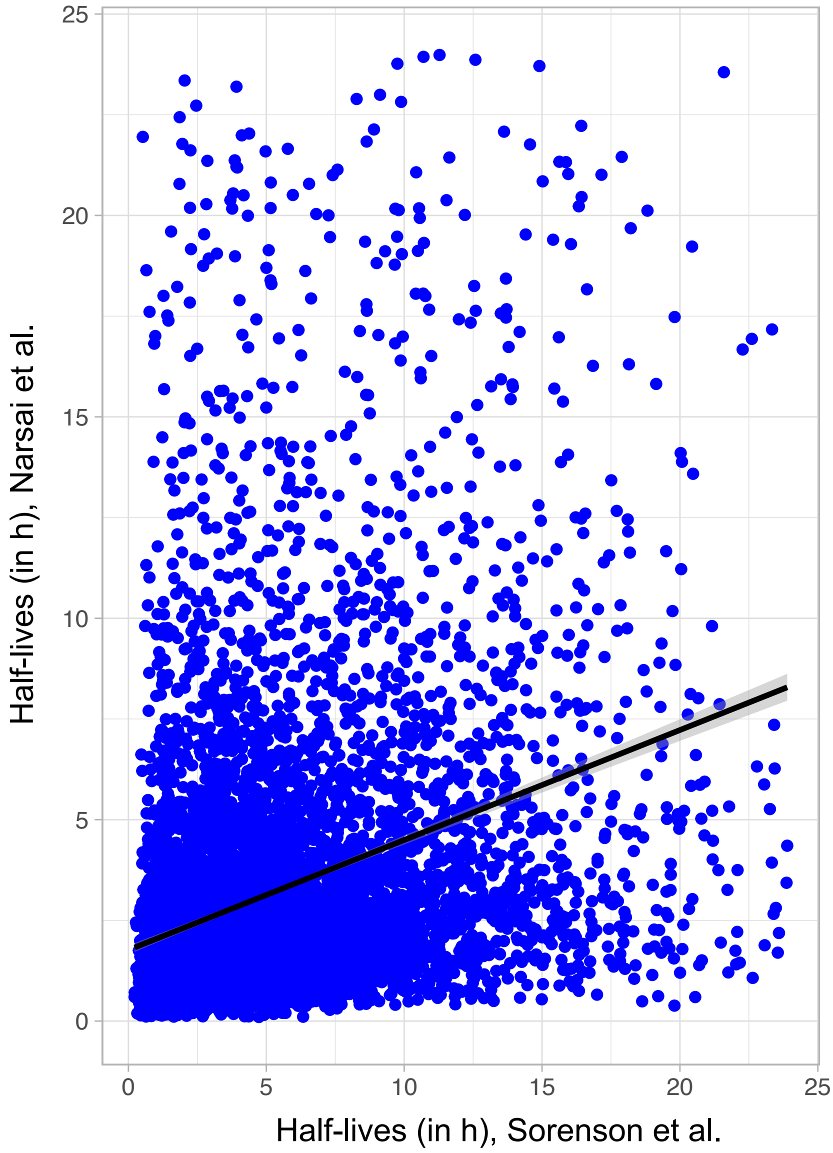


### Figure S3

**Comparison of RNA half-lives estimated by inhibitor-based approaches.**

Comparison of RNA-half-lives determined by Narsai et al. and Sorenson et al.. Narsai et al. applied the transcriptional inhibtor actionmycin D to Arabidopsis cell cultures. Sorenson et al. applied the transcriptional inhibtor cordecypin to Arabidopsis seedlings.
