## Supplementary material for "Metabolic labeling of RNAs uncovers hidden features and dynamics of the *Arabidopsis thaliana* transcriptome": Figure S1

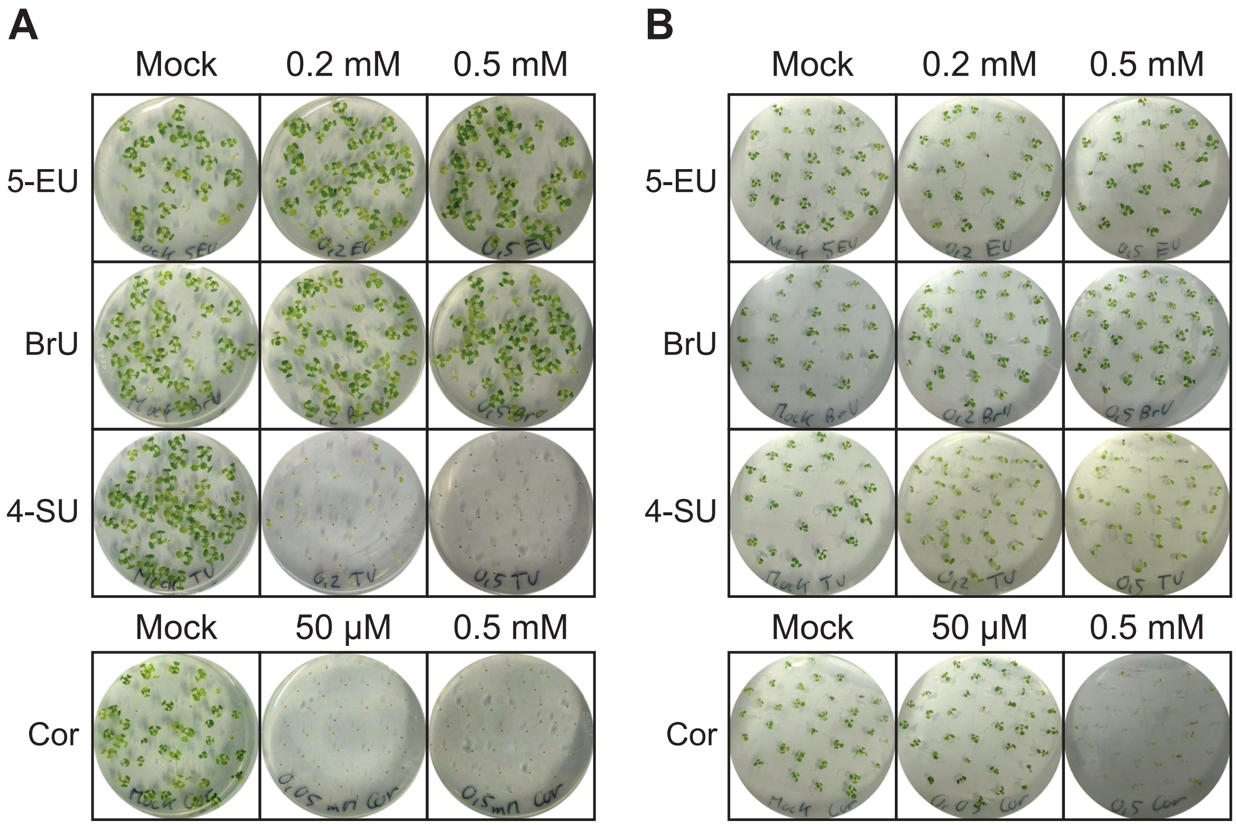


### Figure S1

**Effects of bromouridine (BrU), 5-ethynyl uridine (5-EU), 4-thiouridine (4-SU), and cordycepin (Cor) on plant development**

(A) Arabidopsis plants grew on plant half-strength MS medium containing the indicated amounts of BrU, 5-EU, 4-SU, and cordycepin for 14 days under long-day conditions.

(B) Arabidopsis plants grew for five days on plant half-strength MS medium. Seedling were transferred to half-strength MS medium containing the indicated amounts of BrU, 5-EU, 4-SU, and cordycepin. Seedlings developed for another 14 days under long-day conditions.
