## Supplementary material for "Metabolic labeling of RNAs uncovers hidden features and dynamics of the *Arabidopsis thaliana* transcriptome": Figure S2

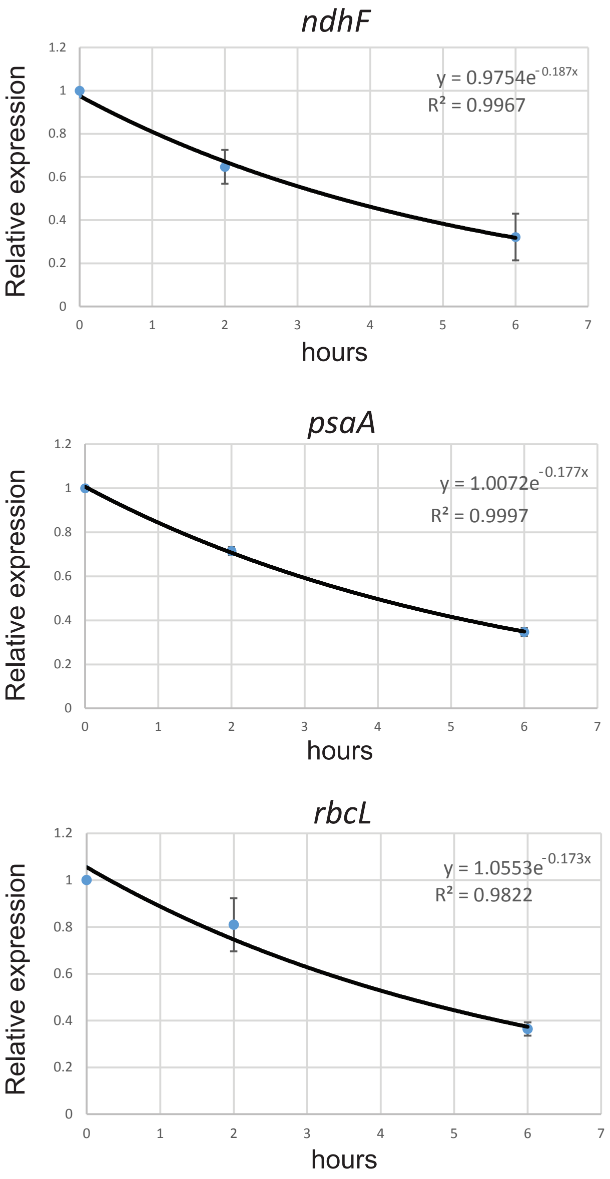


### Figure S2

**Metabolic labeling for RNA stability measurements of chloroplast RNAs**

Decay of selected chloroplast mRNAs measured after 24 hours of 5-EU labelling and subsequent chase with non-labeled uridine. For the analysis of *ndhF* RNA decay, three biological replicates were performed, for *psaA* and *rbcL* two. Graphs represent exponential trend lines with curve equations and R square values displayed in the graph areas.
